# A Dynamic Programming Approach for the Alignment of Molecules

**DOI:** 10.1101/2024.01.23.576849

**Authors:** Ryan A. Williams, David A. Liberles

## Abstract

There is a need for computational approaches to compare molecules based upon chemical similarity or for evaluating biochemical transformations. Here, a new approach to molecular alignment in the Simplified Molecular Input Line Entry System (SMILES) format is introduced. Leveraging dynamic programming and scoring alignments to minimize differences in a measure of electronegativity, the study addresses the challenge of comprehending structural transformations in reaction pathways, notably transitions from linear to cyclical structures. The method is built from the Needleman-Wunsch algorithm for sequence alignment, but with a modified scoring function for different input data. Validation against a benchmarked dataset from the Krebs cycle based upon known chemical mechanisms of molecular transformations confirmed the efficacy of the approach. The transformations of the Pentose Phosphate Pathway were quantified using this algorithm and was compared with Glycolysis. The study introduces an algorithm that quantifies molecular transformations in metabolic pathways, highlighting a midpoint dissimilarity peak in cyclical pathways and a progressive decrease in molecular similarity in linear pathways. The software is available at: https://github.com/rwills5042/SMILES_Alignment.

## II. Introduction

Sequence alignment for biological macromolecules like DNA and proteins is a common precursor to the analysis of these molecules. Sequence analysis is based upon the notion of identifying homology or descent from a common ancestor, but is predicated on the signal for this lying in chemical similarity which is maximized with position-specificity to find an optimal alignment. Except for biochemical transformations, the notion of homology is absent in the alignment of small organic molecules, but the tools for sequence alignment can be reharnessed to align small organic molecules based upon atomic similarity, with position-specific detail. Applications of molecular alignment can include basic questions in biochemistry, including in evolutionary biochemistry, as well as any number of applications in fields from organic synthesis to pharmacology.

Chemical similarity algorithms and programs form the core of various applications such as drug discovery, chemical database management, and structural prediction [1]. Tools like RDKit [2] and Open Babel [3] have provided robust frameworks for generating molecular fingerprints, a popular method for comparing chemical structures. These fingerprints, bit vectors representing the presence or absence of specific structural features, can be compared using metrics such as Tanimoto or Dice coefficients [4], which are simple proportions. Graph-based methods, which treat molecules as graphs with atoms as nodes and bonds as edges, are another common strategy [5]. Algorithms such as the Maximum Common Subgraph (MCS) find the largest shared structure between two molecules [6]. These are approaches for comparing the presence or absence of atoms. In recent years, machine learning methods have emerged as powerful tools for predicting chemical similarity. Neural networks trained to predict molecular properties directly from molecular graphs or SMILES strings have shown promising results [7,8].

Conceptually, molecular comparisons can be bond-centric rather than atom centric, although much of the current focus is on analysis using atoms, sometimes including their neighbors.

Despite these advances, most existing methods do not directly incorporate key physicochemical properties like electronegativity into their similarity measures. Moreover, handling gaps in alignment remains a challenging issue, limiting their utility in dealing with complex molecular structures. This highlights the need for alignment-based algorithms. The more common metric used in bioinformatics and chemoinformatics to calculate the similarity of molecular structures is the Tanimoto coefficient (Tc), which is a simple presence/absence proportion [9].

The Tanimoto Coefficient is simple and can be used to traverse through large databases, however this simplicity is the source of many criticisms. One significant concern is its scale dependency, as it can produce differing results based on the size of the molecules being compared, making it a less ideal choice for contrasting larger molecules with smaller ones [10]. The similarity for smaller structures might be overly amplified [10], although one can conceptually normalize by the number of atoms considered. Furthermore, the coefficient’s binary nature means it doesn’t account for the actual count of a specific feature within a molecule. It considers a feature that appears once and ten times in two different molecules as identical [10]. Despite giving a quantitative measure of similarity, it doesn’t offer a direct interpretation concerning specific chemical attributes, bonds, or structures contributing to the similarity. Also, there is no well-defined rule for setting a threshold for similarity or dissimilarity. The definition of “similar” can vary according to different contexts [11].

Here, we use an electronegativity measure as a core feature of atomic behavior within molecules towards a scoring function to align molecules, establishing correspondences between atoms or groups. By integrating Gasteiger charges into the scoring function, it offers a more detailed perspective on atomic interactions and relationships, making it a valuable addition to the toolkit for understanding and analyzing molecular structures and transformations.

Gasteiger charges, also known as Gasteiger-Marsili charges, are partial atomic charges calculated through an empirical method [12]. These charges approximate the partial charges that would be obtained from a full quantum mechanical treatment of a molecule [12]. The calculation method is based on the electronegativity equalization method (EEM) [12]. The Gasteiger-Marsili partial charge computation returns a set of partial charges for each atom in the molecule, resulting in an output of a list (or other data structure) of these partial charges, one for each atom in the molecule [12].

The Gasteiger charge computation involves an iterative procedure where charges are redistributed among connected atoms in each step, considering their different electronegativity and hardness values [12]. The process of calculating Gasteiger charges begins with an initial estimation of the partial atomic charges based on the atom type, this estimation is typically derived from a lookup table that contains precalculated charges for different atom types. [13]. This initial estimation is based on the principle of electronegativity equalization, an empirical observation that, in a stable molecule, the electronegativity of each atom tends to equalize with that of the surrounding atoms [13]. The method then employs an iterative procedure to adjust the initial charges until the total potential of the molecule is minimized [13]. An iterative method adjusts atomic charges based on electronegativity differences between bonded atoms [13]. For each bond, the charge differences are computed and proportionally applied to the atoms involved, maintaining the molecule’s overall neutrality[13]. This iterative adjustment continues until minimal charge changes are observed, indicating convergence [13]. Subsequently, the calculated charges are normalized to match the molecule’s total charge[12]. Readily available Python modules exist to calculate such charges, such as RDKit [2].

Our approach is centered around the creation of scoring matrices that are based on Gasteiger charges, drawing inspiration from the BLOSUM matrices utilized in protein sequence alignment [14]. We will construct two distinct types of scoring matrices: the first will compare all charges in the metabolism database, while the second will focus on the charges of specific atom pairs such as (C,O), (C,C), and (O,O) within this database. Building upon this groundwork, we will modify the Needleman-Wunsch algorithm, a standard tool in bioinformatics for sequence alignment, to enable the alignment of SMILES molecules using our specially developed scoring matrices [15]. To validate our modified algorithm, we will apply it to a known cyclical biochemical pathway, the Krebs cycle, using a truth set. This application will not only test the algorithm’s precision and reliability but also establish a benchmark for its usage in analyzing more complex molecular pathways. Finally, we will employ the algorithm to examine the Pentose Phosphate Pathway. This step will demonstrate the practical applications of our method in biochemical and molecular research, providing new insights into molecular interactions and alignment within this pathway.

## III. Methods

### IIIa. Database

In this study, focusing on metabolic pathways is of crucial importance as these sequences of reactions form the central framework of biochemistry, offering a rich context to align molecules based on their role in shared or similar chemical reactions. Understanding the similarities and differences between molecules in these pathways can offer valuable insights into metabolic processes and their evolution, facilitating an in-depth analysis of the dynamic nature of metabolic reactions. The alignment of molecules from metabolic pathways can potentially help decipher the molecular basis of transitions between linear and cyclic modes under varying cellular conditions, a critical aspect in predicting metabolic behavior.

The study analyzed molecules from metabolic pathways, using data from PubChem and focusing on the Metabolism Pathway from Reactome (R-HSA-1430728) containing 1101 molecules [16]. To ensure consistency and accuracy in representing these metabolic molecules, each molecule was represented by its SMILES string. The SMILES strings recovered from PubChem were canonized using Indigo python package, ensuring a standardized description of the molecular structures [17].

### IIIb. Canonization

Canonization of SMILES representations is essential for the successful application of alignment algorithms in the comparison of metabolic pathways. In the absence of canonical forms, different representations of the same molecule may yield inconsistent and unreliable alignment scores, complicating the interpretation of results. Canonical SMILES ensure a consistent string representation for the same molecular structure, irrespective of the way it was initially encoded. Blind importation of pathways may be insufficient, proper data processing is necessary.

### IIIc. AllChem Module and Gasteiger charges

There are two scoring matrices were created for the final alignment: an All vs All and paired scoring. All_vs_All_Scoring.py procures SMILES representations of molecular structures of each molecule in the metabolism database, from which it computes the Gasteiger charges, quantities that encapsulate the relative electronegativities of atoms. The obtained charges are aggregated into a master list containing all charges in all molecules. Then, non-finite values (NaN and infinities) are filtered out.

The absolute difference between every possible pair of charges, is determined generating an exhaustive list of charge differences. A probability is determined by dividing the count of charge differences or greater within a given interval by the total number of observations. The score for each interval is then determined by taking the base-2 logarithm of this probability. To achieve a balance between computational efficiency and accuracy, the range of observable charge differences is discretized into intervals of 0.1, spanning from 0 to 3. For each of these discrete intervals, the relative frequency of observed charge differences within that interval is computed. As a result, a scoring matrix is constructed, which provides a quantitative measure of the likelihood of any two atoms having a certain charge difference. The primary objective of the scoring matrix is to incentivize minor electronegativity variations while penalizing significant discrepancies between aligned atoms. To establish a clear benchmark for this, the mean of all computed differences was ascertained. This average, approximately 0.3, serves as the threshold in the scoring matrix: differences exceeding this average are met with penalties, while those below it is awarded favorable scores.

Paired_Scoring.py is dedicated to the generation of a paired scoring matrix. The scoring function within this structure is based on the differences of Gasteiger charges between paired atoms. The objective is to uncover any underlying trends or patterns exclusive to these specific atomic pairs, such as all Carbon-Carbon (C-C), Carbon-Oxygen (C-O), etc., across all molecules in the database. By comparing these charge differences across a wide range of molecules, the study aims to identify potential trends or phenomena that might enhance our understanding of the behavior of these atoms within the realm of metabolic reactions that the All vs. All scoring matrix is not capable of. For each atom pair, a scoring matrix similar to that in All_vs_All_Scoring.py is created, with the difference that it bases its scores on the specific pair’s charge differences.

### IIId. Algorithm

All_vs_All_Alignment.py focuses on the sequence alignment of two molecules represented in the SMILES format by use of a modified Needleman Wunsch alignment algorithm [15]. Before aligning the molecules, the script processes the SMILES by stripping the characters, and then aligns based on the electronegativity pattern of the molecule. Once the alignment is achieved, the original characters are replaced. This is done so that non-characterizable elements (bonds, charges, and rings) do not interfere with the alignment. The align() function in All_vs_All_Alignment.py is the modified Needleman-Wunsch alignment algorithm. When the scoring matrices are being filled, the scores are filled in the traditional manner, however the scores are determined by the difference of charge of the two atoms using the designed scoring matrix. Align() returns the two aligned molecules and an alignment score. Paired_Alignment.py carries a notable distinction from All_vs_All_Alignment.py as it utilizes the scoring matrix devised in Paired_Scoring.py.

### IIIe. Pathways

The Krebs cycle, also known as the citric acid cycle or tricarboxylic acid cycle, was chosen for the validation of the algorithm due to several reasons. Primarily, it is one of the most well-studied and understood biochemical pathways in cellular metabolism. This high level of understanding ensures that the true transformations and mechanisms involved in each step are well-documented, providing a solid ground truth against which to compare the algorithm’s predictions. Furthermore, the Krebs cycle is relatively short with eight distinct enzymatic steps, simplifying the process of validation [18]. Each step in the cycle involves the transformation of one molecule into another, which aligns well with the purpose of the algorithm, to compare and align molecular structures [18]. Moreover, the cyclical nature of the Krebs cycle provides additional benefits for validation. In a cycle, the product of the final step is the substrate for the first step [18]. This allows for a unique validation perspective, not only can individual transformation accuracy be identified but also the ability of the algorithm to correctly align and track transformations across the entire cycle. Overall, the combination of the Krebs cycle’s simplicity, cyclical nature, and well-established scientific understanding makes it an ideal choice for validating the accuracy and utility of the molecular alignment algorithm. It can also be used to generate data to address evolutionary questions about the evolution of new enzymatic steps and the degree of chemical transformation that typically happens in a step, including questions of if and how linear pathways can become circular ones or about the retrograde hypothesis for pathway evolution [19].

In the approach, coenzymes were excluded from the analysis. This decision was driven by the role coenzymes typically play in the metabolic process. Often acting as electron or functional group carriers, coenzymes do not structurally transform in the same way primary metabolites do [18]. Their inclusion could introduce unnecessary complexity, potentially obscuring the transformation patterns we’re interested in studying.

To ensure consistency and maintain accurate representation of the Krebs cycle environment, a standardized representation was made where all molecules were depicted with carboxylate groups (for example) using the Indigo toolkit [17]. In the final set of SMILES strings, the order of atoms, the identification of functional groups, and carbon chains were clearly discernible [17].

### IIIf. Validation

The validation of the alignment algorithms was conducted using a carefully curated truth dataset, centered on the Krebs cycle. The Krebs cycle, integral to the metabolic functions of aerobic organisms, was streamlined by removing coenzymes and focusing exclusively on the seven primary metabolites [18]. Each metabolite was aligned with the other metabolites. The purpose of aligning all molecules with each other is to validate the algorithms’ ability to differentiate changes each molecule makes as it progresses through the metabolic cycle. An exhaustive analysis of the seven metabolites’ structure and composition was undertaken, with crucial carbons and functional groups identified [18]. Figure 3 shows an example of the validated alignment process of the first metabolite Oxaloacetate aligned with the other metabolites, where the left shows the entire SMILES aligned and the right shows what the algorithm will interpret. It is worth noting that for each glucose molecule metabolized, the citric acid cycle completes two revolutions [18]. Nevertheless, for the purpose of simplicity and to maintain a clear focus on the fundamental transformations, the validated alignment was restricted to the first cycle only. The alignment on the left in Figure 3 is one with the characters in the SMILES intact, while the alignment on the right is the alignment with the characters removed. While the input of the align function is the SMILES with the characters, the alignment itself is done based on the electronegativity blueprints where the non-atomic characters are stripped. The alignment itself will be compared to those on the right set of alignments in Figure 3.

The validation of the optimal parameters involved a comprehensive examination of the alignment parameters - gap open and gap extension penalties. The range for these parameters was chosen between −5 and +5 to ensure a broad coverage of possible scenarios and enhance the robustness of the optimal parameters. Every possible combination of these parameters was tested. This approach was designed to identify the most fitting set of parameters that yielded the highest alignment scores and best represented the structural and functional similarities and dissimilarities between the metabolites in the Krebs cycle. Based on existing knowledge and the fundamental principles of sequence alignment, certain expectations were put forward [15]. The gap open penalty was anticipated to be more penalizing than the gap extension penalty for similar alignments. This is based on the understanding that introducing a gap in the alignment is a more significant event and should be penalized more heavily than simply extending an already opened gap [15]. The comprehensive testing allowed for a rigorous examination of these expectations and served as a robust strategy for the identification of the most fitting parameters for this particular alignment problem. The parameters identified as optimal were those that satisfied these theoretical expectations and also performed best in the practical task of accurately aligning the metabolites in the Krebs cycle.

The optimal parameters were determined by using two methods of similarity: Levenshtein similarity and an exact measurement, while aligning the validated alignments and the experimental alignments [20].

## IV. Results

### IVa. All vs All Scoring and Alignment

Figure 1 shows the plotted All vs. All scoring Matrix. As the observed difference increases, the calculated score decreases, indicating a lesser probability of encountering large charge differences during molecular alignment. Interestingly, the slope of the graph is significantly steeper within the range of 0-1.2, reflecting a more rapid decline in score as the observed difference increases within this interval. Beyond 1.2, up to 3, the slope becomes less steep, indicating a more gradual decrease in score with increasing observed difference. This inflection at 1.2 could suggest that smaller charge differences (less than 1.2) are more common in the dataset, while larger differences (greater than 1.2) occur less frequently.

**Figure 1:**
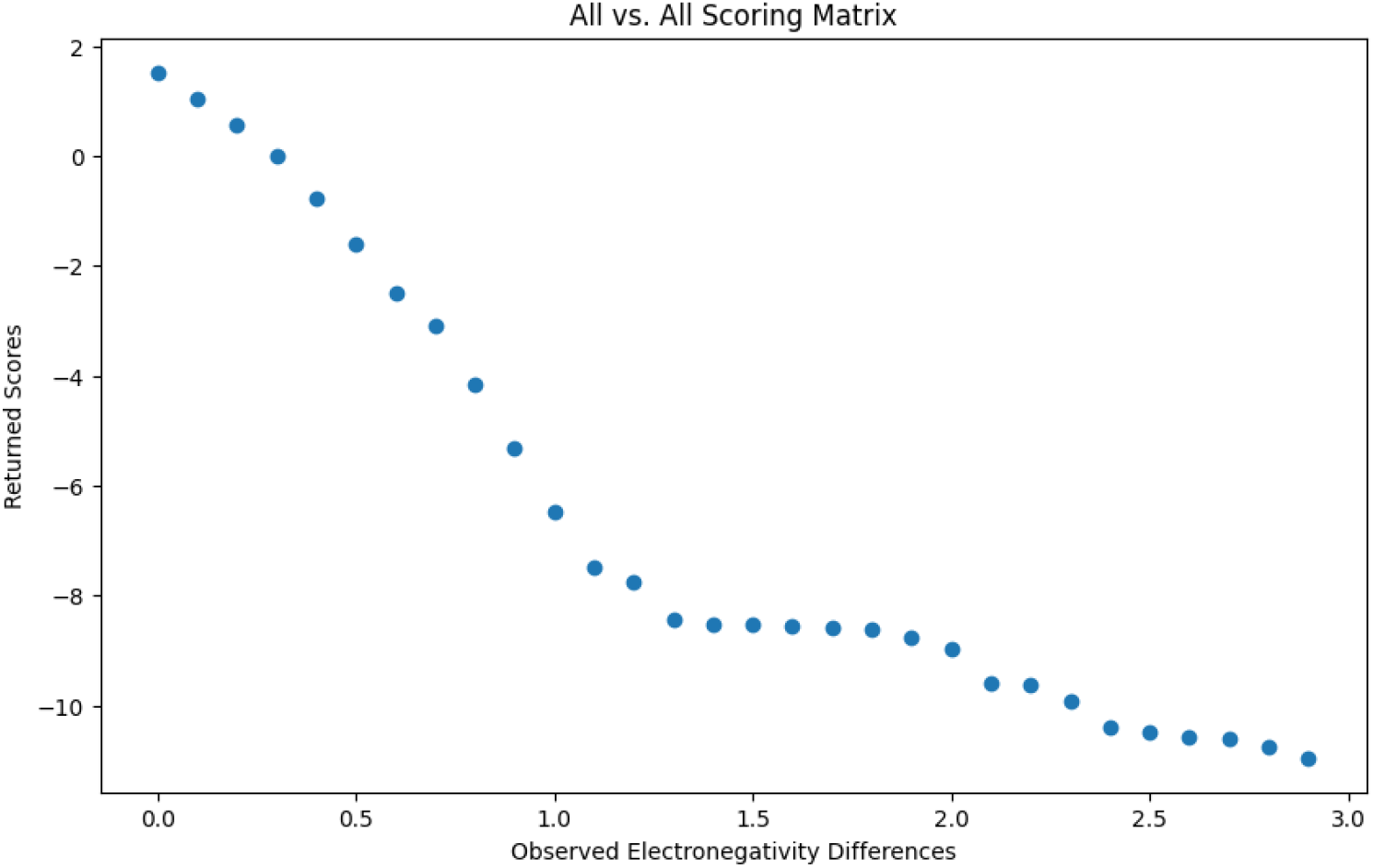
All vs. All scoring Matrix Plot. Log2 probability of observed Gasteiger charge differences or greater in 0.1 increments.

**Figure 2:**
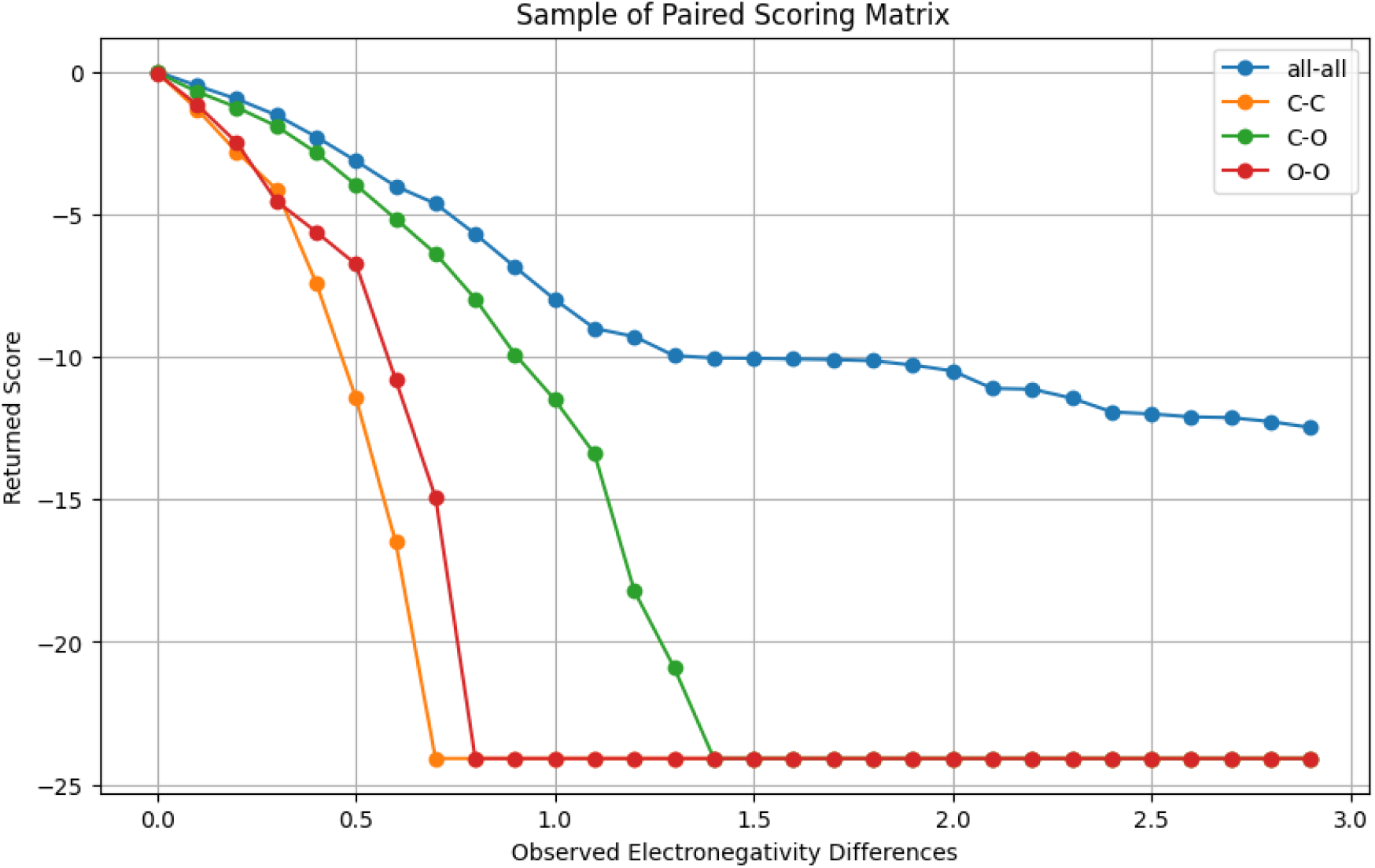
Common Pairs in the Paired Scoring Matrix and All vs. All Scoring Matrix. Log2 probability of observed Gasteiger charge differences or greater in 0.1 increments of each atom pair in the database.

**Figure 3:**
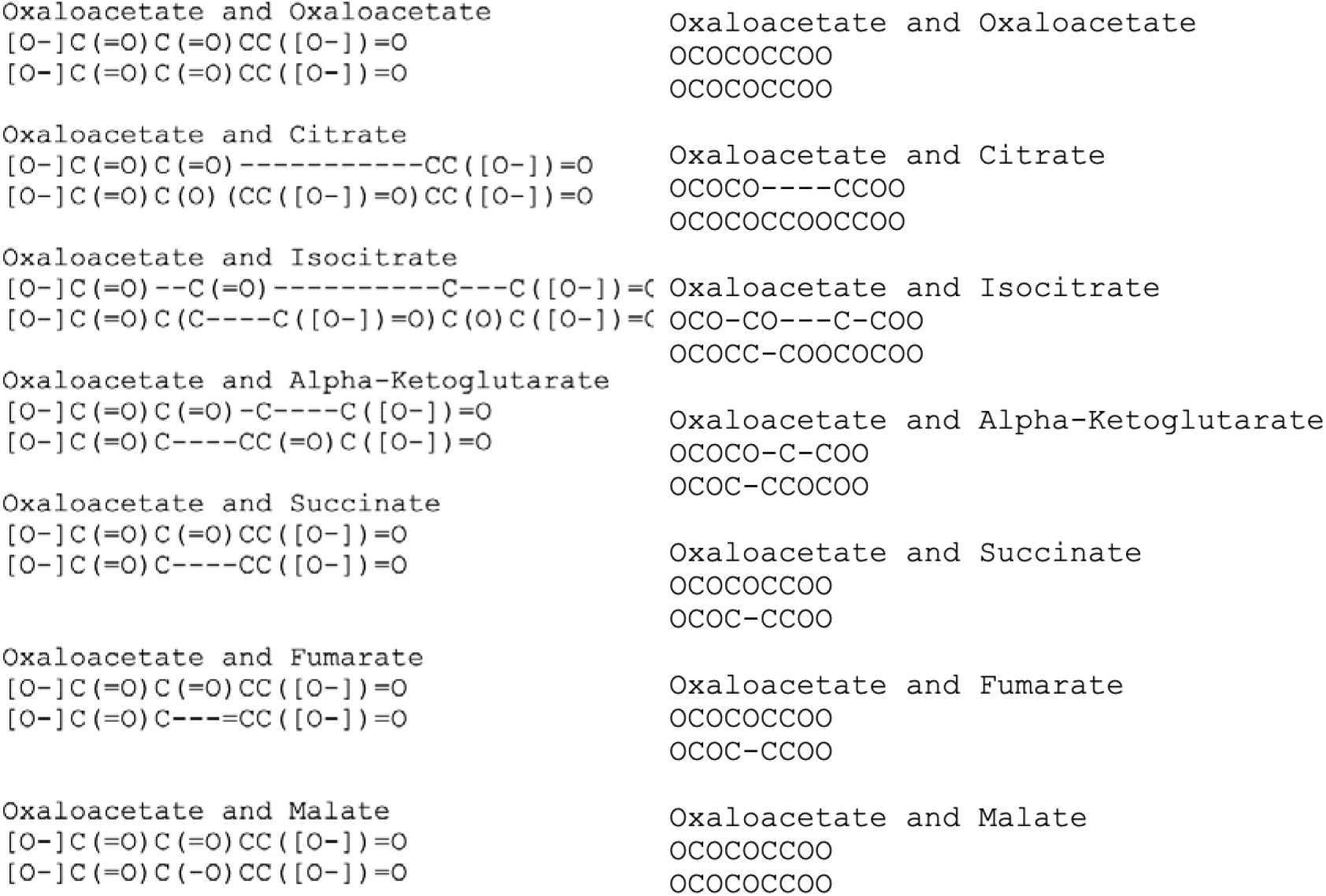
Sample of the Truth Dataset of the Krebs cycle Alignment With Oxaloacetate, with SMILES characters (Left) and without SMILES characters (Right). Left figure demonstrates the truth data set used in the validation, the right figure demonstrates what the algorithm will interpret.

Table 1 presents the top 10 outcomes derived from the all-versus-all scoring matrix for both Levenshtein (Left) and Exact (Right) string similarity. Although not in the same order, both tables have the same gap open and gap extension combinations. A standout observation from this table is the consistently high similarity average in both Levenshtein and Exact, which surpassed 0.95 and 0.94 respectively, underlining a marked degree of similarity amongst the aligned sequences.

**Table 1:** Validation of the All vs All Scoring Matrix on Krebs cycle Truth Dataset for Levenshtein (Left) and Exact (Right). The All vs All scoring alignment algorithm was validated using the truth dataset and determined using Gap open and gap extension penalties ranging from −5 to +5. The accuracy of the alignment was measured using Levenshtein similarity and exact similarity.

|  | Gap Open Penalty | Gap Extend Penalty | Levenshtein Average | Exact Average |  | Gap Open Penalty | Gap Extend Penalty | Levenshtein Average | Exact Average |
| --- | --- | --- | --- | --- | --- | --- | --- | --- | --- |
| 32 | -2 | 0 | 0.958939 | 0.908744 | 45 | 0 | 0 | 0.955228 | 0.940019 |
| 45 | 0 | 0 | 0.955228 | 0.940019 | 39 | -1 | 0 | 0.951518 | 0.910600 |
| 39 | -1 | 0 | 0.951518 | 0.910600 | 32 | -2 | 0 | 0.958939 | 0.908744 |
| 0 | -5 | -5 | 0.942680 | 0.883739 | 0 | -5 | -5 | 0.942680 | 0.883739 |
| 24 | -3 | 0 | 0.939540 | 0.880599 | 24 | -3 | 0 | 0.939540 | 0.880599 |
| 22 | -3 | -2 | 0.939540 | 0.880599 | 22 | -3 | -2 | 0.939540 | 0.880599 |
| 21 | -3 | -3 | 0.939540 | 0.880599 | 21 | -3 | -3 | 0.939540 | 0.880599 |
| 31 | -2 | -1 | 0.939540 | 0.880599 | 31 | -2 | -1 | 0.939540 | 0.880599 |
| 15 | -4 | 0 | 0.939540 | 0.880599 | 15 | -4 | 0 | 0.939540 | 0.880599 |
| 14 | -4 | -1 | 0.939540 | 0.880599 | 14 | -4 | -1 | 0.939540 | 0.880599 |

Figure 4 visualizes the interplay between gap open parameters and the resulting similarity average from the alignments for Levenshtein and Exact. A notable trend is observed: as the gap open penalty becomes less penalizing (increases numerically), there is a corresponding decrease in the similarity average. This trend is intuitive when considering the principles of sequence alignment. A reduced gap open penalty means that the algorithm is more permissive or lenient towards introducing gaps into the alignment. Consequently, this leniency can lead the algorithm to frequently insert gaps in the alignment rather than matching or mismatching characters, even if such a choice may not reflect the genuine similarities between sequences. When gaps are preferentially favored over matches or mismatches due to a lessened penalty, the resulting alignments tend to exhibit decreased similarity values. This underscores the importance of judiciously setting the gap open penalties: while making them too harsh can deter necessary gaps, making them too lenient can lead to overuse of gaps, compromising the genuine representation of similarities in the aligned sequences.

**Figure 4:**
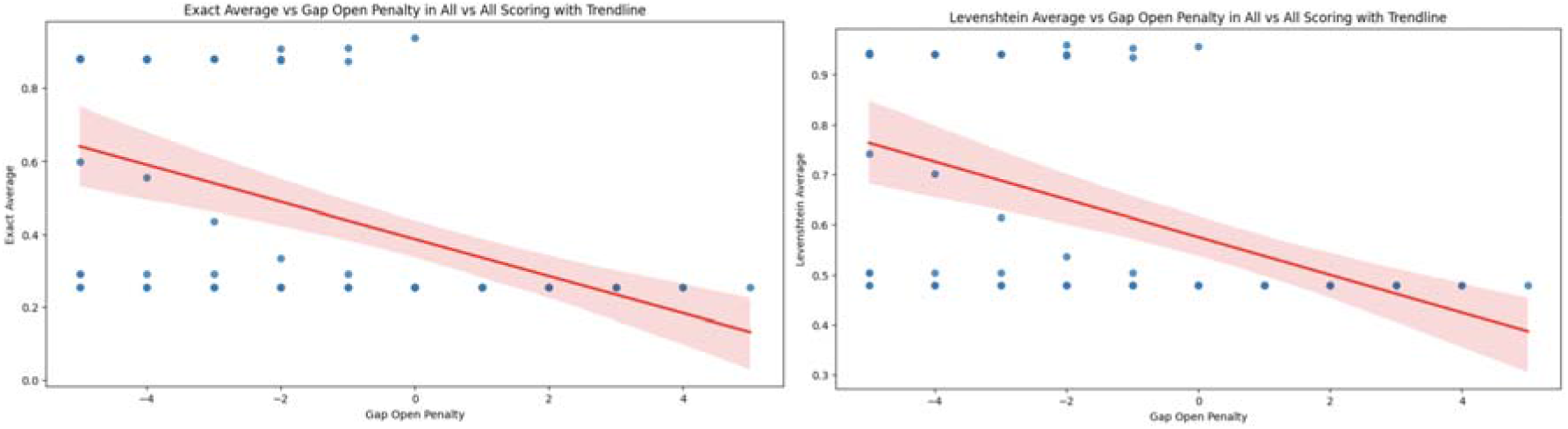
Similarity Average vs Gap Open Penalty in All vs All scoring for Levenshtein (Left) and Exact (Right). As the gap open penalty becomes less penalizing (increasing numerically), there is a corresponding decrease in the similarity average in the All vs All scoring alignment algorithm regardless of accuracy measurement.

Figure 5 presents the relationship between gap extension penalties and the similarity average derived from the alignments for Levenshtein and Exact. A discernible trend emerges: as the gap extension penalty becomes less penalizing (increasing numerically), the similarity average also decreases. This trend aligns with our understanding of the alignment process. When the penalty for extending an already existing gap is reduced, the alignment algorithm tends to elongate gaps rather than aligning characters from the sequences. By doing so, while it might optimize for reduced penalty costs, it can potentially misrepresent the true alignment or similarity between sequences. Prolonged gaps, driven by a minimal extension penalty, can overshadow the genuine shared patterns between sequences, thus leading to a reduction in the observed similarity values. This observation emphasizes the need for carefully calibrated gap extension penalties: they should neither be too stringent, preventing necessary gap extensions, nor too lenient, encouraging overly extended gaps that could distort the true sequence similarities. Both graphs have discernable drop off once the penalty is greater than 0 and no longer penalizing, which falls in line with the expectations of the algorithm.

**Figure 5:**
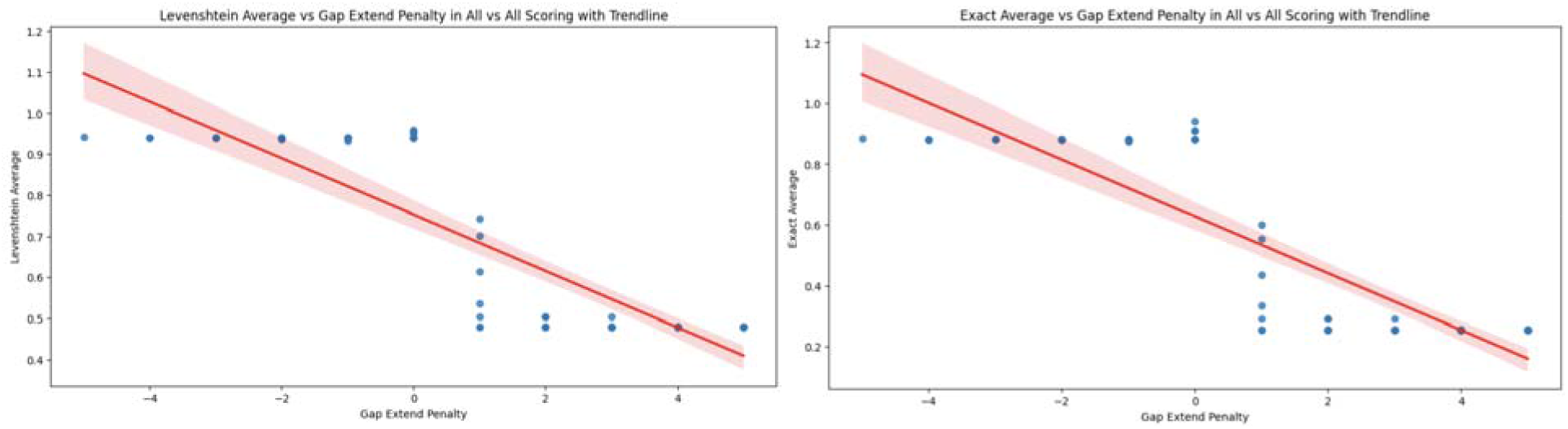
Similarity Average vs Gap Extension Penalty in All vs All scoring for Levenshtein (Left) and Exact (Right). As the gap extension penalty becomes less penalizing (increasing numerically), there is a corresponding decrease in the similarity average in the All vs All scoring alignment algorithm regardless of accuracy measurement with a sharp decrease when the penalty is greater than 0.

Figure 6 presents the relationship between the Exact and Levenshtein similarities in the validation. The positive correlation demonstrates that both methods report similar findings without having bias.

**Figure 6:**
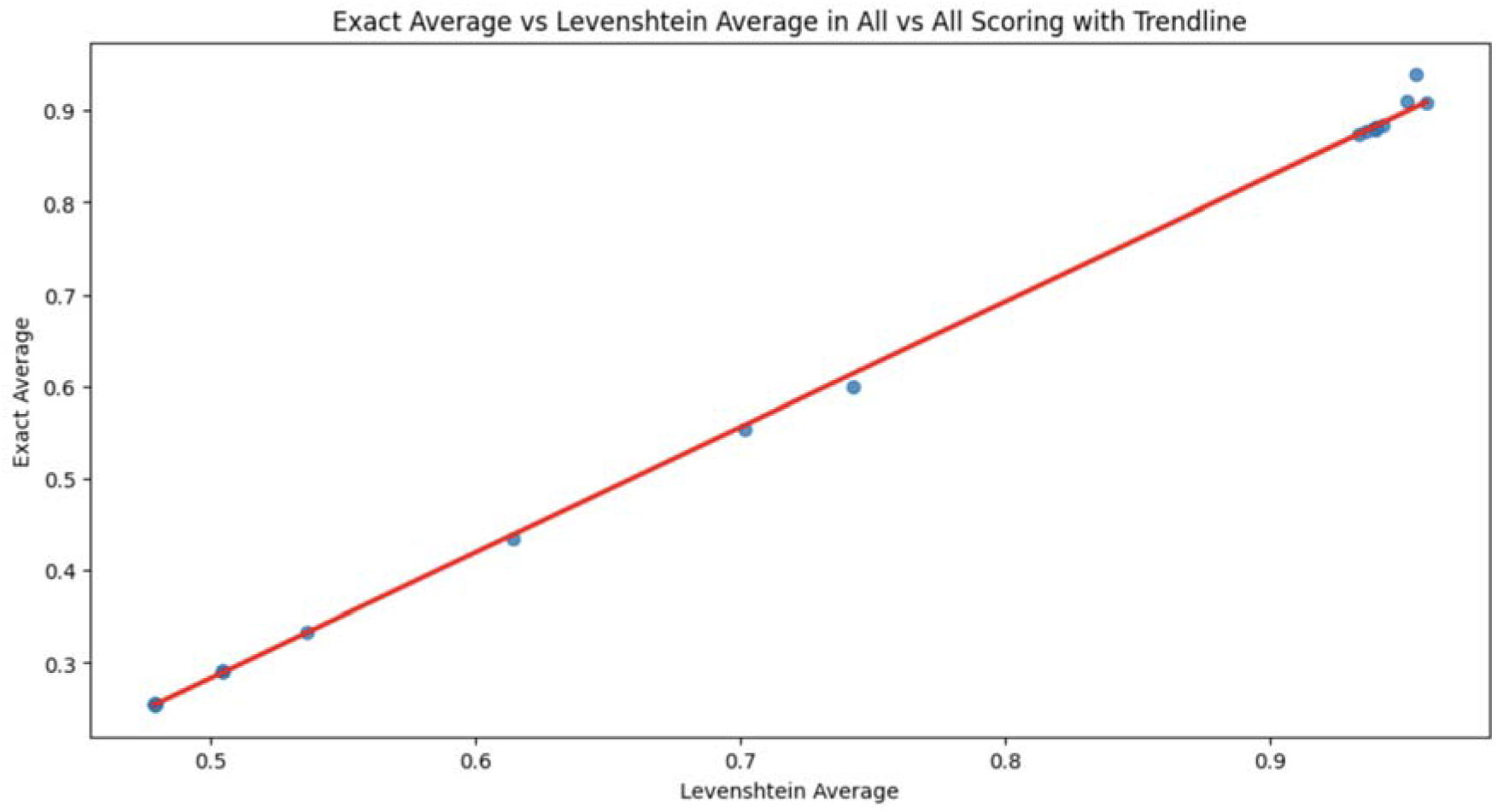
Levenshtein vs Exact Averages in All vs All Scoring. Figure demonstrates no appreciable difference between the two accuracy measurements giving no bias or preference between the two.

### IVb. Paired Scoring and Alignment

Figure 2 presents a graphical comparison between the scoring matrices for the most frequently observed atom pairs (’C’, ‘C’), (’C’, ‘O’), and (’O’, ‘O’) and the all-vs-all scoring matrix derived in the earlier section. The plots for the atom pairs exhibit an inverse quadratic shape initially, suggesting a nonlinear correlation between charge differences and their corresponding scores for small charge differences. However, these plots transition into a line once the scoring matrix reaches its largest penalty, which appears to be at a score of around −15. This shift reflects the enforcement of the largest penalty for extreme charge differences. In contrast, the all-vs-all scoring matrix plot maintains a different profile. It displays an initial steep linear decline that flattens out for larger charge differences, with the maximum penalty positioned approximately at −12.5. This divergence from the atom pair-specific plots underscores the distinctive characteristics of the all-vs-all scoring matrix, reflecting a broader spectrum of atom types and charge differences.

Table 2 presents the top 10 outcomes derived from the paired scoring matrix for both Levenshtein (Left) and Exact (Right) string similarity. Both tables share most gap open and gap extension combinations, once again demonstrating similar responses in similarity methods.

**Table 2:**
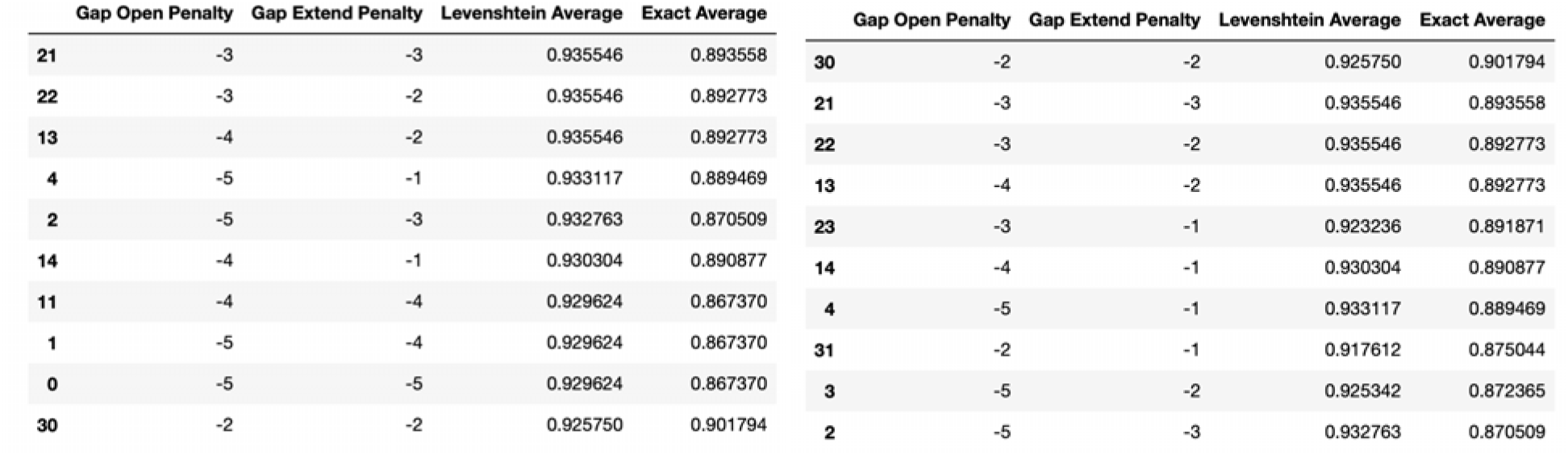
Validation of the Paired Scoring Matrix on Krebs cycle Truth Dataset for Levenshtein (Left) and Exact (Right). The All vs All scoring alignment algorithm was validated using the truth dataset and determined using Gap open and gap extension penalties ranging from −5 to +5. The accuracy of the alignment was measured using Levenshtein similarity and exact similarity.

Figures 7, 8, and 9 exhibit trends analogous to those observed in Figures 4, 5, and 6. Figure 7 mirrors the relationship presented in Figure 4, demonstrating how gap open parameters influence the resulting similarity averages in both Levenshtein and Exact measurements. The discernible trend suggests that as the gap open penalty becomes progressively less stringent, similarity averages correspondingly decline. This behavior is consistent with the understanding that a lower penalty allows the algorithm to frequently introduce gaps, potentially misrepresenting genuine sequence similarities. Figure 8, akin to Figure 5, maps out the correlation between gap extension penalties and the derived similarity averages. The observed trend is clear: easing the gap extension penalties, by increasing their values, results in diminishing similarity averages. This behavior is indicative of the alignment algorithm’s propensity to favor prolonged gaps, especially when the penalties for such extensions are minimized. This might inadvertently lead to overshadowing authentic sequence similarities, necessitating a balanced penalty setting to accurately capture the inherent sequence relationships. Lastly, Figure 9 parallels Figure 6, representing the correlation between the Exact and Levenshtein similarity metrics within the context of the validation using the paired scoring method. The evident positive correlation reaffirms the notion that both metrics yield comparable outcomes, ensuring unbiased representation and validation of string similarities. Despite the distinct scoring methods between Figures 4-6 and Figures 7-9, the trends and insights derived remain strikingly consistent, emphasizing the robustness and reproducibility of the observed relationships across different scoring paradigms.

**Figure 7:**
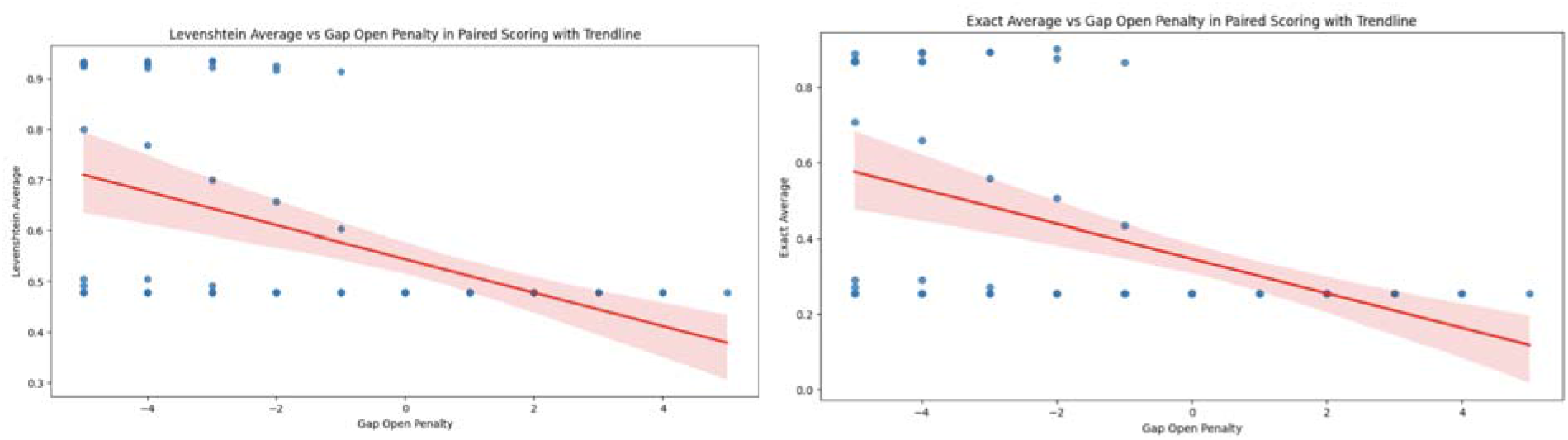
Similarity Average vs Gap Open Penalty in Paired Scoring for Levenshtein (Left) and Exact (Right). As the gap open penalty becomes less penalizing (increasing numerically), there is a corresponding decrease in the similarity average in the All vs All scoring alignment algorithm regardless of accuracy measurement.

**Figure 8:**
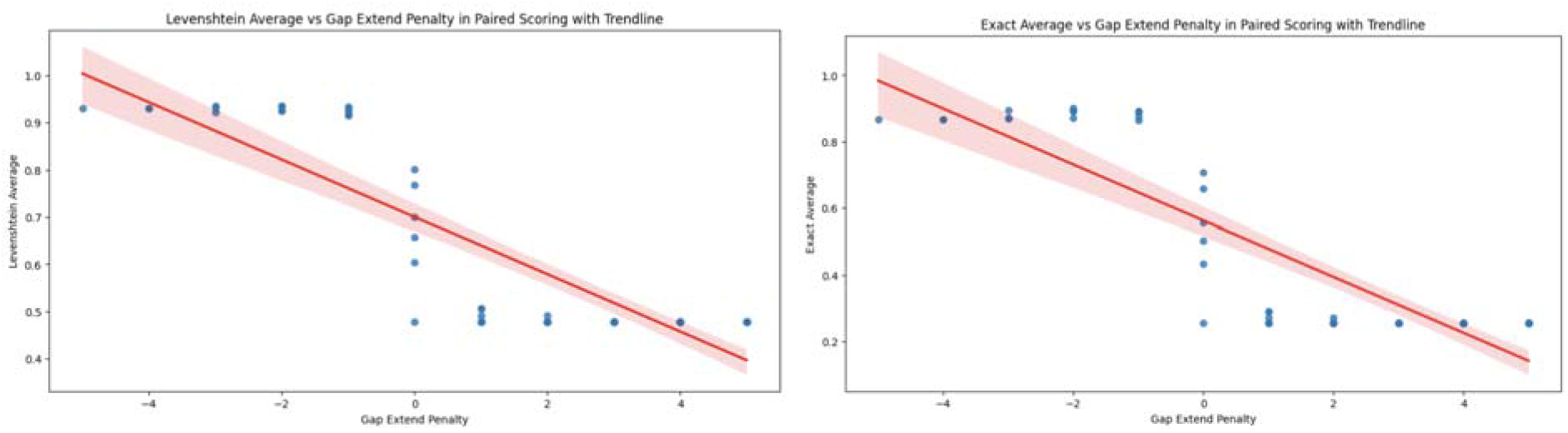
Similarity Average vs Gap Extension Penalty in Paired scoring for Levenshtein (Left) and Exact (Right). As the gap extension penalty becomes less penalizing (increasing numerically), there is a corresponding decrease in the similarity average in the All vs All scoring alignment algorithm regardless of accuracy measurement with a sharp decrease when the penalty is greater than 0.

**Figure 9:**
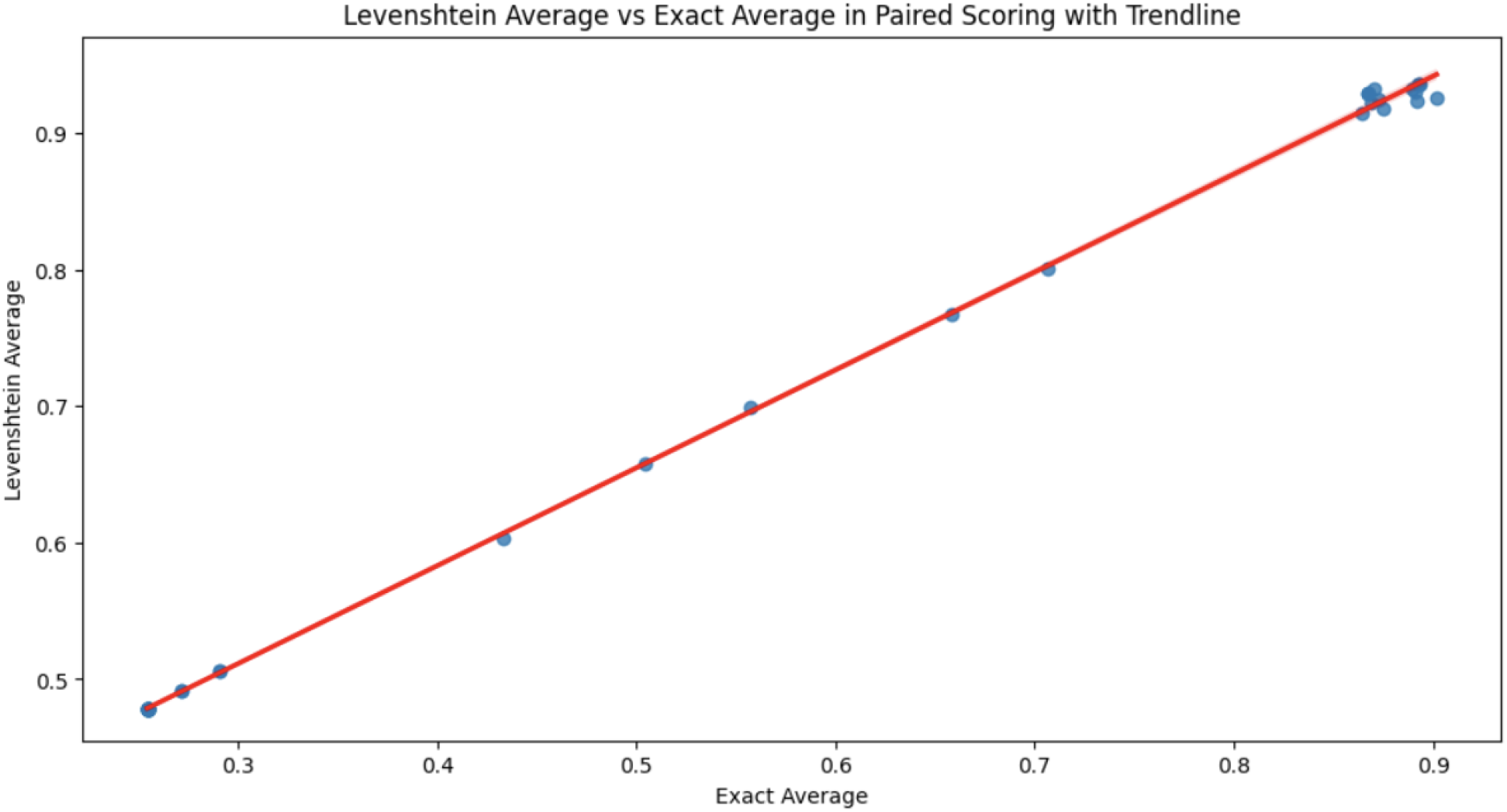
Levenshtein vs Exact Averages in Paired Scoring. Figure demonstrates no appreciable difference between the two accuracy measurements giving no bias or preference between the two.

### IVc. Application

Two prominent models, the retrograde and patchwork evolutionary models, offer distinct perspectives on the evolution of metabolic pathways [19, 21, 22]. The retrograde model posits that metabolic pathways adapted to the scarcity of specific substrates by developing new enzymes for more abundant, upstream compounds [19, 21]. In contrast, the patchwork model suggests that early enzymes with broad substrate specificity gradually specialized through processes like gene duplication and sequence divergence [19, 22]. For some chemical processes, including those that generate ATP, linear pathways can become cyclical ones under some selective constraints, but also with chemical constraints as well.

In the evolution from linear to cyclical pathways, a key characteristic is that molecular transformations do not alter the molecule so drastically that it cannot revert to its original structure; such an extreme change would result in the pathway remaining linear. In a cyclical molecular pathway, how significant can the chemical transformations of any starting molecule be at each step before it returns to its original position? Molecules become increasingly dissimilar from their initial structure up to a certain point in the cycle, after which they begin to revert. The algorithm developed quantifies these transformations, allowing for the tracking of molecular changes, pinpointing key transformation steps, and identifying when molecules start resembling their original state. Furthermore, it facilitates comparative analysis across different pathways, potentially uncovering patterns in the evolutionary shift from linear to cyclical pathways.

The Pentose Phosphate Pathway (PPP) is a vital metabolic route that functions parallel to glycolysis. It chiefly assists in the generation of ribose-5-phosphate, which is vital for nucleotide synthesis, and in the production of NADPH, an indispensable cofactor for reductive biosynthesis reactions within cells [23]. Utilizing both the all vs all and paired scoring algorithms on this pathway, as retrieved from the KEGG database [24], provides insights into molecular transformations throughout the PPP.

Figures 10 and 11, demonstrates the average alignment score for each minimum distance in the cycle for the best parameters found in the validation for each step in the pathway, for both All vs All and Paired Scoring methods respectively. Both figures follow the expectations of a cyclical cycle similarity: descending in similarity, reaching a point of most dissimilar, and approaching more similar as the transformations return to its starting position. For both scoring methods, the most dissimilar positions appear most distant from a given position. For comparison, the Krebs cycle was analyzed using the same parameters (Figures 12 and 13), and demonstrates the expectation described above and that is seen in the PPP. The Krebs cycle alignments also show that the most dissimilar part in the cycle is also the most distant position. The error bars show high variance for both figures, likely due to the different sizes of the molecules which have large impacts on the alignment scores. It is possible for large pathways that molecules will asymptote out at a high level of distance.

**Figure 10:**
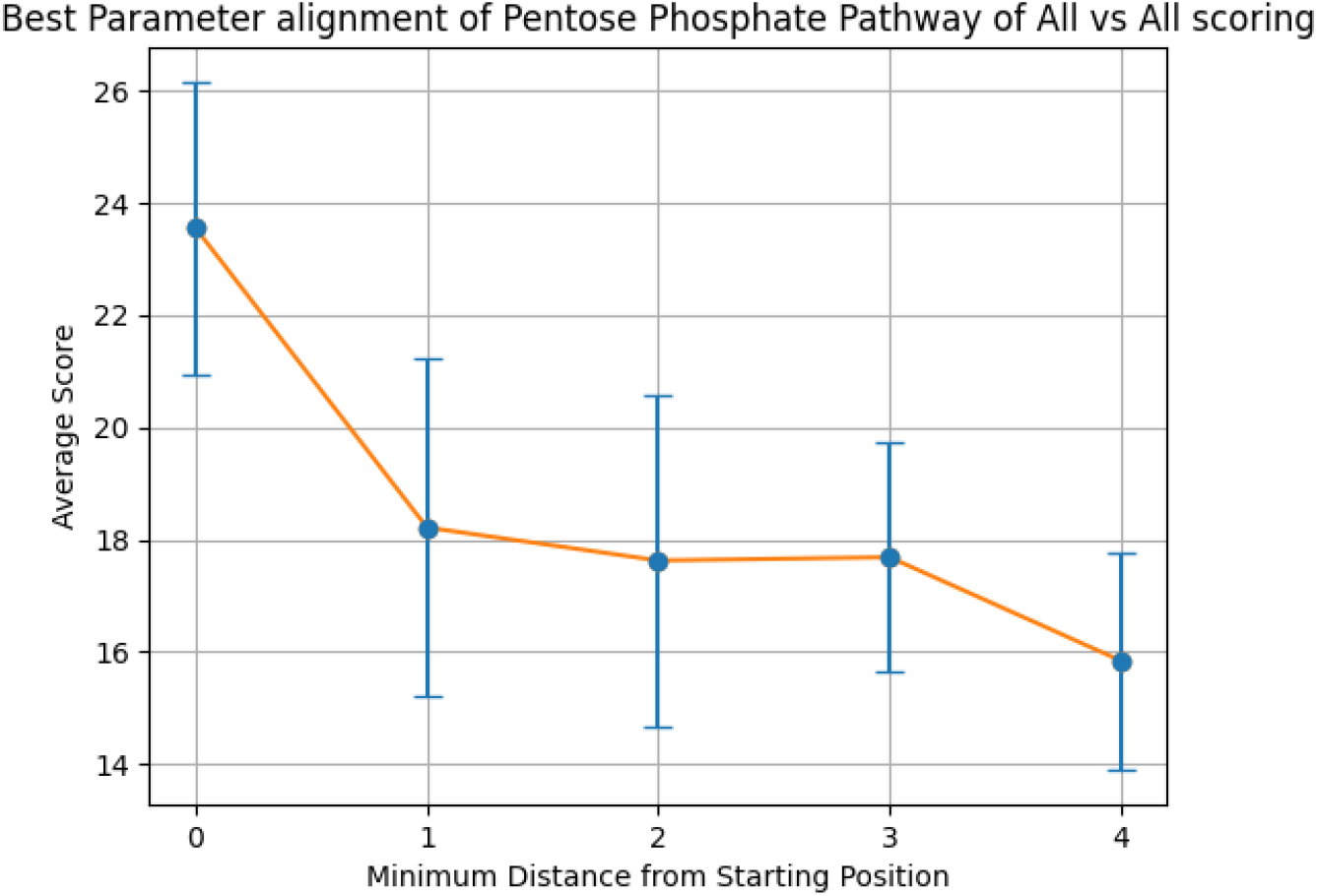
Alignment of the PPP using all vs all scoring matrix. Using the best parameters (0 for both gap open and gap extension determined in the validation step), each step of the cyclical pathway was aligned with the subsequent steps so that the entire cycle is aligned for each step. The y axis shows the average score for each distance from the starting position (x axis). Standard Deviations [2.60195146, 3.01405173, 2.95498863, 2.03380675, 1.93686312].

**Figure 11:**
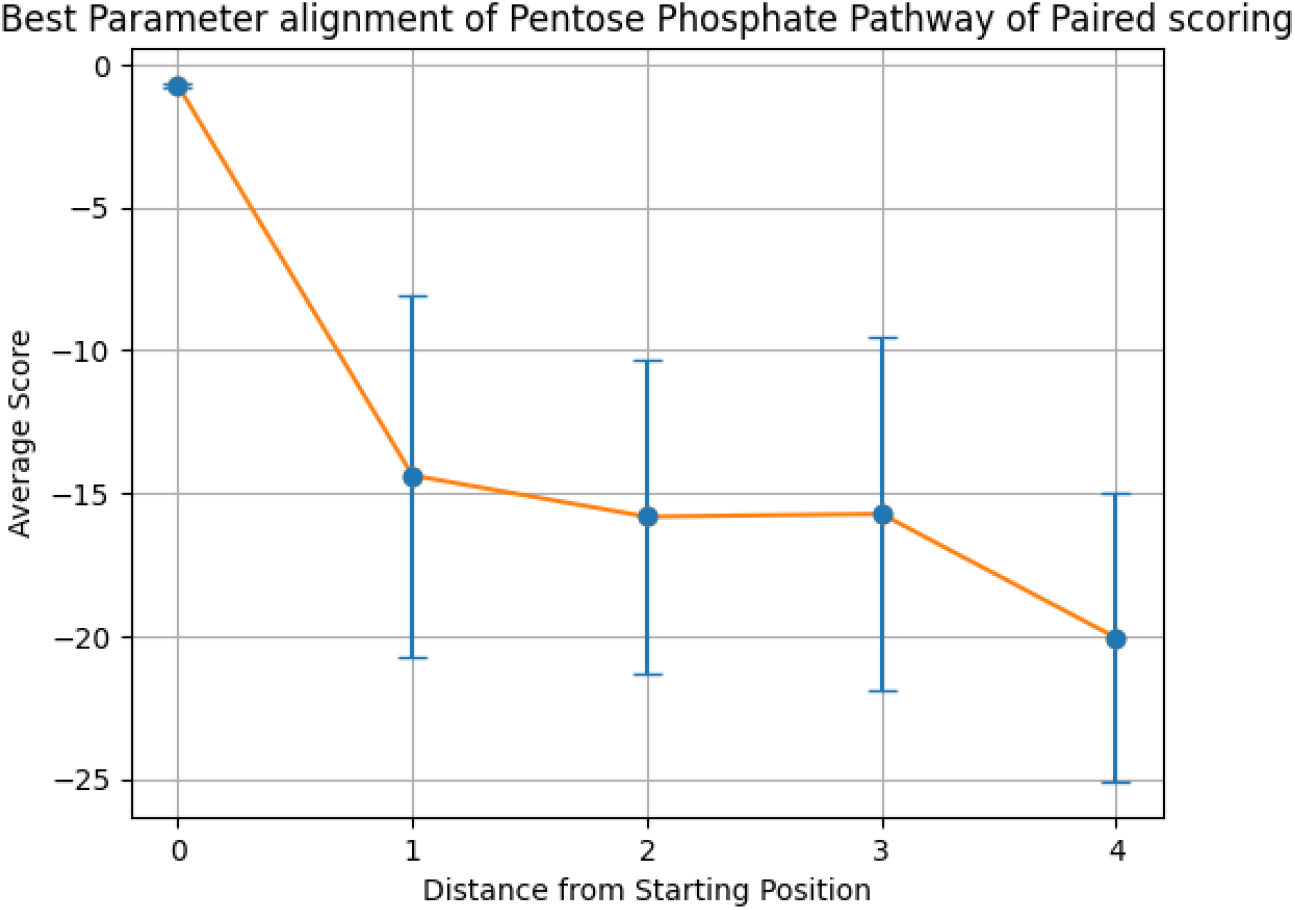
Alignment of the PPP using paired scoring matrix. Using the best parameters (−2 for both gap open and gap extension determined in the validation step), each step of the cyclical pathway was aligned with the subsequent steps so that the entire cycle is aligned for each step. The y axis shows the average score for each distance from the starting position (x axis). Standard Deviations: [0.07839836, 6.33330154, 5.51964565, 6.20545232, 5.036306]

**Figure 12:**
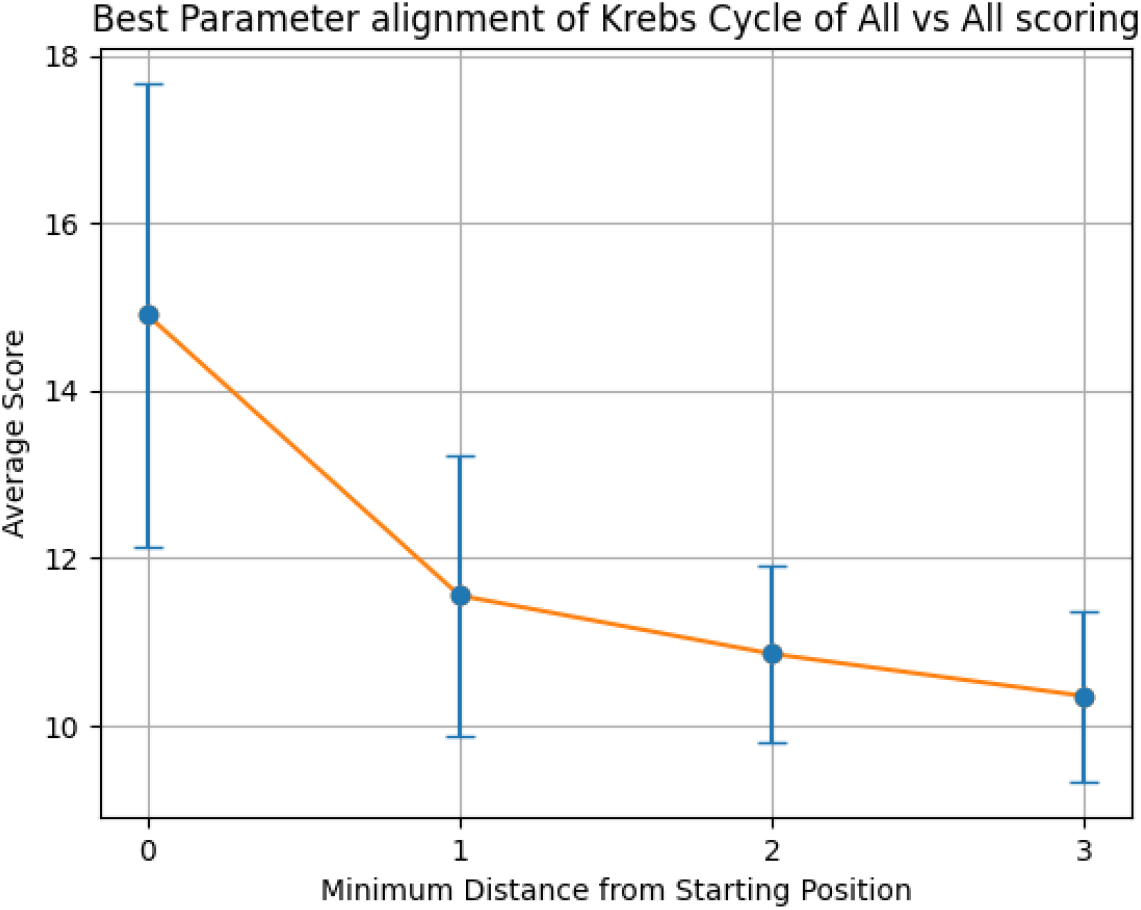
Alignment of the Krebs cycle using all vs all scoring matrix. Using the best parameters (0 for both gap open and gap extension determined in the validation step), each step of the cyclical pathway was aligned with the subsequent steps so that the entire cycle is aligned for each step. The y axis shows the average score for each distance from the starting position (x axis). Standard Deviations: [2.76238373, 1.68189704, 1.05823526, 1.02090974]

**Figure 13:**
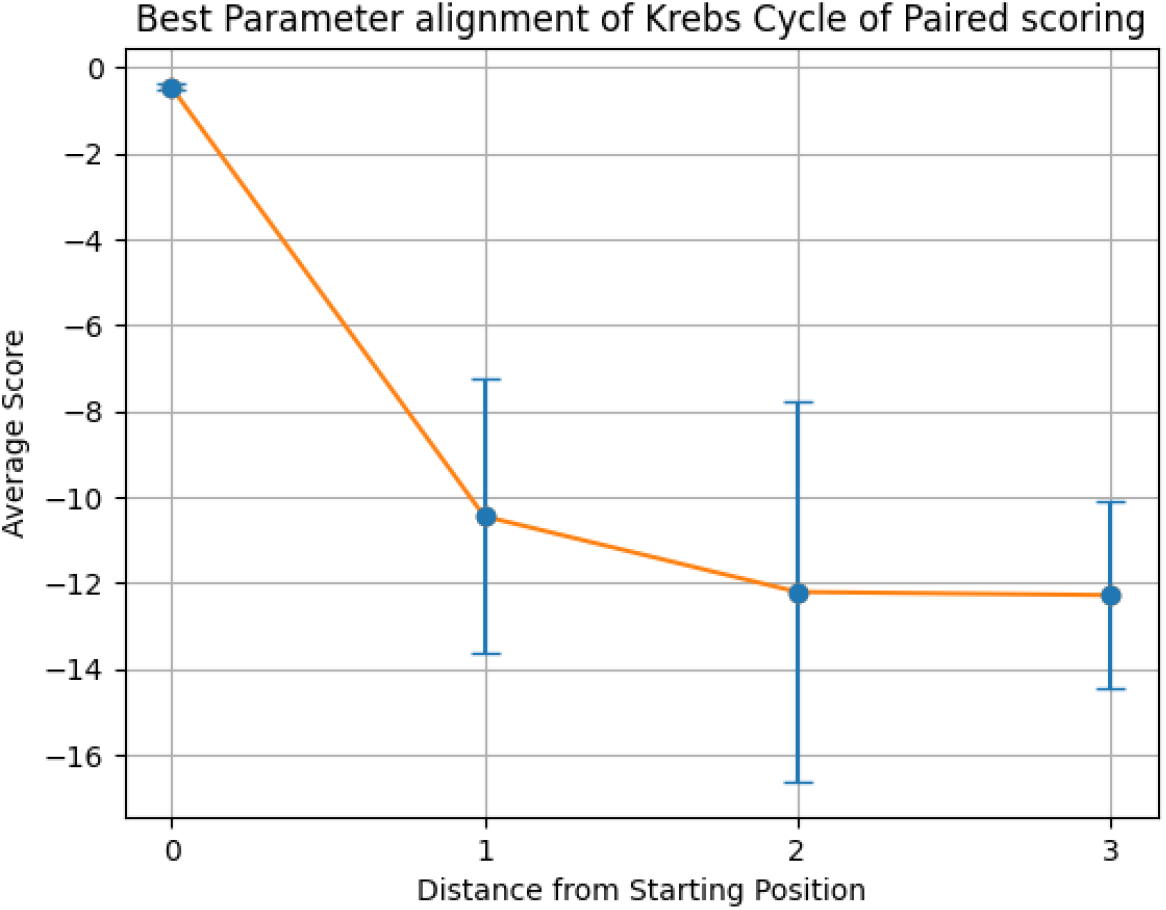
Alignment of the Krebs cycle using paired scoring matrix. Using the best parameters (−2 for both gap open and gap extension determined in the validation step), each step of the cyclical pathway was aligned with the subsequent steps so that the entire cycle is aligned for each step. The y axis shows the average score for each distance from the starting position (x axis). Standard Deviations: [0.08527199, 3.18849359, 4.41807495, 2.19116527]

Figures 14 and 15 display the alignment scores for the Glycolysis pathway using All vs All and Paired Scoring methods, respectively. Consistent with the expected behavior of a linear pathway, the figures illustrate an increase in dissimilarity as the distance from the starting molecule grows. This trend in the alignment scores reaffirms the algorithm’s validity, complementing the patterns observed in Figures 10-13 for cyclical pathways, where dissimilarity peaks midway and then decreases. Collectively, these results validate the algorithm’s efficacy in capturing the anticipated divergence patterns in both linear and cyclical metabolic pathways.

**Figure 14:**
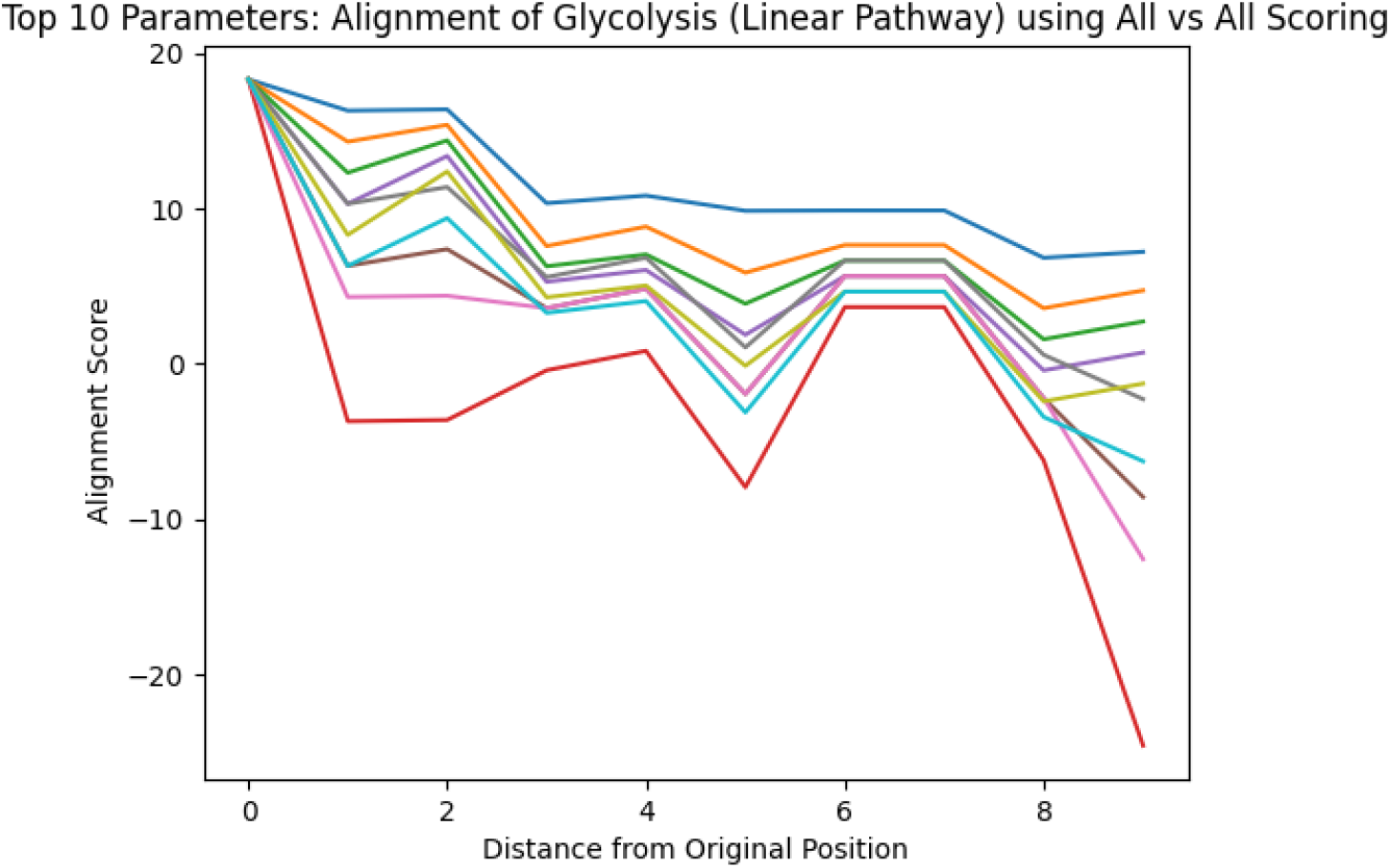
Alignment of Glycolysis Pathway using All vs All Scoring. Using the top 10 parameters, the linear pathway glycolysis was aligned using the All vs All scoring algorithm, demonstrating the algorithms ability to show decreasing alignment scores expected in linear pathways.

**Figure 15:**
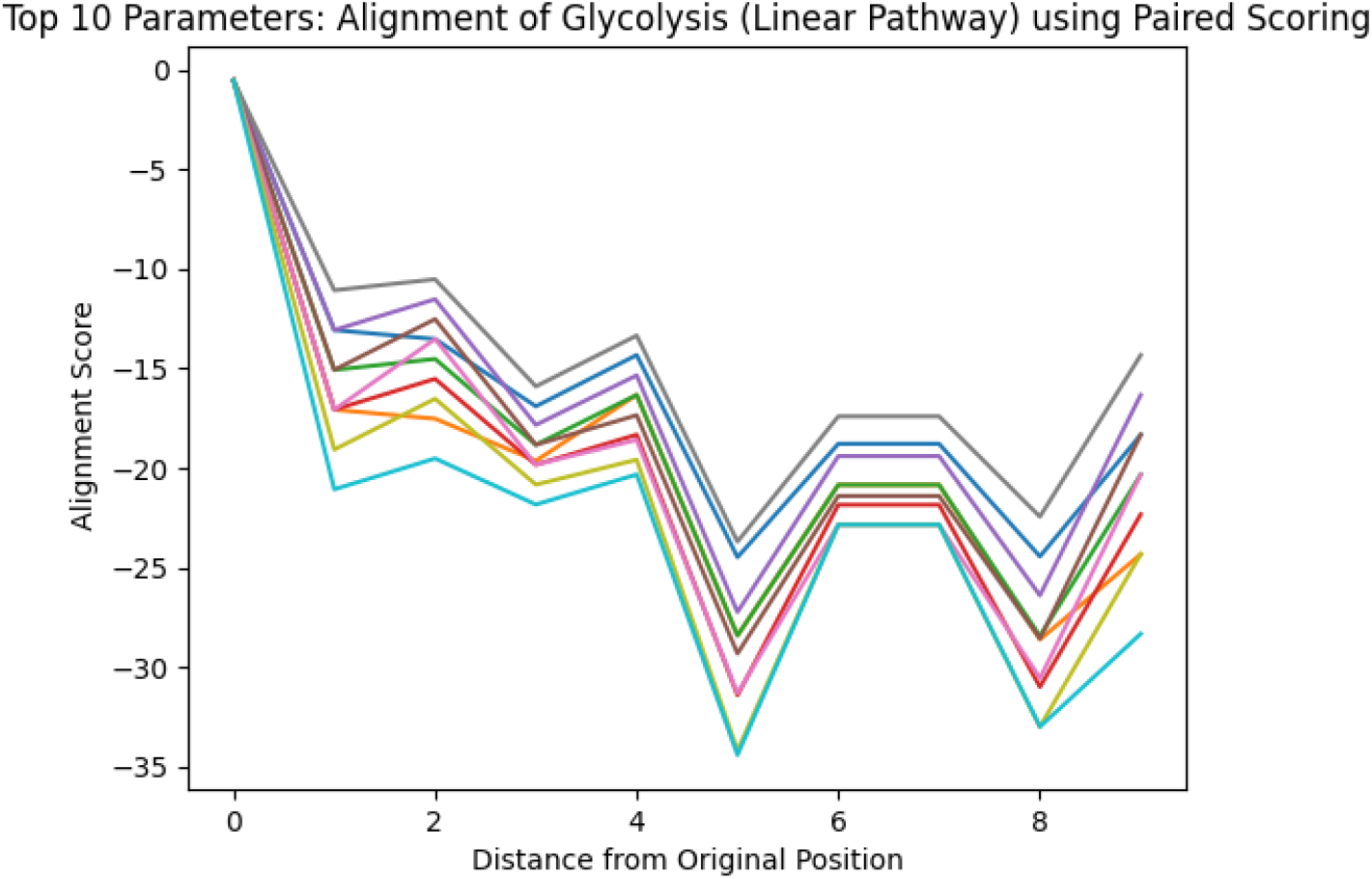
Alignment of Glycolysis Pathway using Paired Scoring. Using the top 10 parameters, the linear pathway glycolysis was aligned using the Paired scoring algorithm, demonstrating the algorithms ability to show decreasing alignment scores expected in linear pathways.

## V. Discussion and Conclusion

The original Needleman-Wunsch algorithm, being a well-established method in the field of bioinformatics, has been made widely available in various scientific programming libraries [15]. Its general simplicity, combined with the documentation available, ensured that its integration into the pipeline was significantly expedited. The development of the custom SMILES alignment algorithm, seen as an expansion on the Needleman-Wunsch, was facilitated efficiently due to its foundational reliance on an existing approach [15].

In the alignment process for molecules represented as SMILES, the decision to strip non-atomic characters was driven by the need to distill the essence of the molecule to its electronegativity blueprint. SMILES notations are rich in detail, encompassing both atomic and non-atomic characters. While this offers a comprehensive representation, it introduces the challenge of aligning non-characterizable entities, which would introduce unnecessary noise during the alignment process. By eliminating these characters, the focus shifts entirely to the alignment of the underlying electronegativity patterns intrinsic to each atom. It’s pertinent to note that while explicit characters indicating certain molecular features are absent post-stripping, the retained electronegativity is not an isolated characteristic; it’s deeply influenced by both the atom type, bond type, and its spatial orientation within the molecule. Thus, the alignment process, by focusing on this electronegativity blueprint, effectively captures the core nature and orientation of atoms within molecules, ensuring a more refined and accurate alignment devoid of the potential distractions introduced by non-atomic characters.

The research and subsequent results on the alignment algorithms notably bridge a gap in the fields of chemoinformatics and bioinformatics. While both domains have seen rapid advancements, the specific challenge of molecular alignment, especially as it pertains to intricate biochemical pathways like the Krebs cycle, has largely remained a complex puzzle. This study’s achievements offer a tailored solution, catering not only to the general alignment needs but also to the subtleties and nuances of biochemical transformations. By building upon and refining established alignment techniques, this tool provides an optimized approach, meticulously adapted to chemical structural data.

The algorithm has unveiled that within cyclical metabolic pathways, the point of greatest dissimilarity from the starting molecule typically occurs halfway through the cycle. This discovery suggests a threshold of molecular alteration beyond which a linear pathway may not naturally evolve into a cyclical one. Moreover, the analysis indicates that in linear pathways, similarity diminishes as the distance from the starting molecule increases, implying an inverse relationship between similarity and pathway progression. These findings could help in predicting the evolutionary potential of metabolic pathways, providing a metric to gauge the likelihood of a pathway remaining linear or becoming cyclical based on its molecular transformation scores.

For instance, how might a particular molecule evolve within a metabolic pathway? How can we predict structural transformations in newly discovered or lesser-known pathways? What are the likely structural consequences of introducing a novel compound into a system? The algorithm has the potential to systematically analyze molecular libraries, highlighting evolutionary trends and pinpointing the evolutionary shifts in metabolism. Using large libraries, it can be determined what the threshold for similarity is for cyclical pathways. Furthermore, its predictive power can be leveraged to infer the metabolic roles of molecules that are not yet understood, by evaluating their similarities within the extensive network of metabolic reactions.. Such capabilities could significantly enhance our comprehension of metabolic evolution and the intricate design of biological systems.

## Author Contributions

RAW executed the work and wrote the initial version of the manuscript. DAL conceived and supervised the project and contributed to writing of advanced versions of the manuscript.

## Notes

The authors declare no competing financial interest.

## Acknowledgments

The authors would like to thank Joel Sheffield and Taha Iqbal for discussions early in the generation of this work.

## Data and Software Availability

The software is available at: https://github.com/rwills5042/SMILES_Alignment.

